# vToxiNet: a biologically constrained deep learning framework for interpretable prediction of drug-induced hepatotoxicity

**DOI:** 10.64898/2026.02.26.708259

**Authors:** Xuelian Jia, Tong Wang, Daniel P. Russo, Lauren M. Aleksunes, Shuo Xiao, Hao Zhu

**Author notes:** X.J. and T.W. contributed equally to this work.

## Abstract

Hepatotoxicity remains a leading cause of drug attrition and post-marketing withdrawal, resulting from diverse and complex toxicity mechanisms. Traditional *in vitro* models can only capture a limited subset of toxicity pathways, and animal studies face translational and ethical limitations. Regulatory agencies have therefore promoted new approach methodologies, including human-relevant assays, omics technologies, and computational models to improve predictive toxicology and support evidence-based decision-making. However, most machine learning models for hepatotoxicity either rely solely on chemical structure or operate as black boxes, limiting mechanistic interpretability and broader applicability. Here, we introduce the virtual toxicity network (vToxiNet), a biologically constrained deep learning framework that embeds systems toxicology knowledge directly into neural network architecture for interpretable hepatotoxicity prediction. vToxiNet integrates chemical descriptors, high-throughput assay responses, transcriptomic signatures, and Reactome pathway hierarchy to construct a virtual adverse outcome pathway network. Across cross-validation and multiple external validation datasets, vToxiNet demonstrates robust predictive performance and generalizes to previously unseen chemicals. Importantly, interpretation of vToxiNet enables gene and pathway-level attribution, supporting mechanism-informed hazard characterization and chemical prioritization. These results demonstrate that encoding biological hierarchy as architectural constraints enables both predictive accuracy and mechanistic insight, establishing a generalizable framework for modeling complex biological outcomes.

## 1. Introduction

The liver plays a central role in metabolism and detoxification of xenobiotics, thus is vulnerable to injury by potential toxic chemicals including drugs. Hepatotoxicity remains a major challenge in drug development, often leading to the failure of promising candidates in clinical trials and serving as the most common cause of market withdrawal due to adverse effects after approval^1–4^. Hepatotoxicity is influenced by various toxicity pathways, reflecting intricate toxicity mechanisms as the drugs interact with biological systems in diverse ways^2^. Regulatory agencies require a standard battery of tests to evaluate the safety and toxicity of drugs and chemicals in humans, as outlined in the International Council for Harmonisation of Technical Requirements for Pharmaceuticals for Human Use (ICH) and Organisation of Economic Cooperation and Development (OECD) guidelines. This framework relies heavily on animal studies, which require significant time and resources, and remain insufficient to address the growing need for efficient toxicological evaluations of diverse drug candidates and environmental chemicals^4, 5^. Moreover, translating animal study results to humans remains a significant challenge, particularly in cases of idiosyncratic hepatotoxicity, due to species differences in metabolism and physiology as well as the complexity of underlying mechanisms^2, 6^. A retrospective analysis revealed that animal testing missed 45% of hepatotoxicity cases during clinical trials^7^. This highlights the urgent need for alternative testing strategies that are more predictive of human outcomes.

Over the past few decades, advances in molecular biology, high-throughput screening (HTS) and machine learning approaches have enabled the development of alternative strategies for chemical toxicity evaluation, such as using human-relevant assays and computational modeling. These alternative strategies, often referred to as new approach methodologies (NAMs), aim to reduce reliance on animal testing while improving human relevance to support predictive toxicology and regulatory decision-making^8, 9^. The rapidly growing volume of HTS and toxicogenomics data generated by NAMs has fueled the growth of toxicity big data, providing a strong foundation for machine learning–based toxicity modeling and prediction.

However, most existing machine learning models for hepatotoxicity, such as quantitative structure-activity relationship (QSAR) models, were primarily developed using only chemical structure information^10–14^ and therefore do not fully exploit the wealth of big data now available for chemical risk assessments. Furthermore, widely used machine learning approaches, such as deep neural networks, often produce “black box” models that lack transparency in their predictive processes and do not meet international guidelines that emphasize mechanistic interpretation of toxicity^15, 16^. To resolve this issue, incorporating a mechanistic knowledge framework, such as adverse outcome pathways (AOPs) and systems toxicology, into model design can enhance the interpretability of model predictions and provide toxicity insights to support decision-making^17–19^.

In previous studies, we developed a predictive AOP model that assessed hepatotoxic liability for new toxicants through forming reactive metabolites leading to oxidative stress and subsequent hepatotoxicity^20, 21^. However, hepatotoxicity can also arise from other mechanisms, including organelle dysfunction, altered liver metabolism, and immune-mediated responses.

Therefore, the reported oxidative stress related model can only reveal a portion of hepatotoxicants that induce toxicity through this specific mechanism. To pursue a global model that can cover multiple hepatotoxicity mechanisms, we developed the virtual toxicity network (vToxiNet), an innovative biologically constrained deep neural network (DNN) approach designed to enhance the predictability and mechanistic understanding of hepatotoxicity (**Fig. 1A**). vToxiNet leverages the Reactome pathway ontology to construct a virtual AOP network encompassing a wide spectrum of toxicity pathways. This knowledge-based modeling framework integrates multi-modal data for model training, including the chemical structure information, HTS assay responses, gene expression profiles and gene-pathway relationships representing potential key events in hepatotoxicity. As a global hepatotoxicity modeling tool, vToxiNet can assess the liabilities of new drug molecules by providing mechanistic insights into the contributions of different pathways to the cellular responses that may culminate in liver injury.

**Fig. 1.**
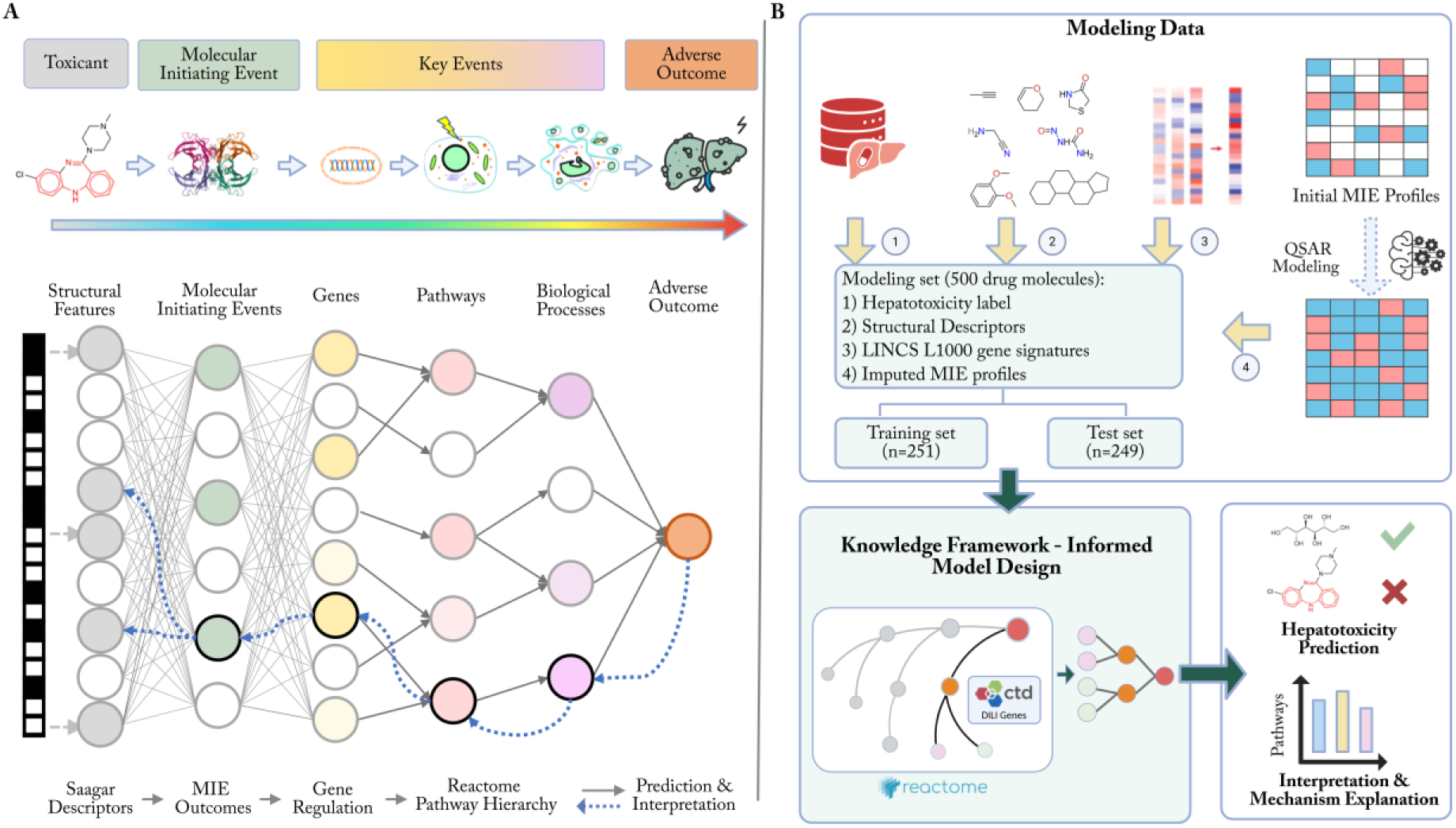
Interpretable knowledge-informed deep learning modeling framework. **(A)** Schematic representation of vToxiNet architecture. The design of vToxiNet follows the general AOP framework, where a toxicant binds to a receptor (MIE), triggering changes in gene expression and subsequent molecular and cellular key events, ultimately leading to an adverse outcome. The vToxiNet architecture mirrors the AOP framework by encoding different biological entities into a deep neural network with customized, biologically informed connections. **(B)** Modeling data and knowledge integration for the design and training of vToxiNet. The knowledge framework, including the Reactome pathway hierarchy and CTD curated gene-disease interactions, was incorporated into the model design. Multimodal data including hepatotoxicity labels, chemical descriptors, MIE assay responses, and gene expression data, were integrated for model training. Finally, model predictions and interpretation were used to infer mechanistic explanations.

## 2. Results

### 2.1. Overview of datasets

The multimodal data used for vToxiNet modeling included hepatotoxicity labels, structural descriptors, MIE assay responses, and L1000 gene expression profiles (**Fig. 1B**). Saagar fingerprints were calculated and used as structural descriptors, while other datasets were compiled from public sources. The hepatotoxicity dataset was primarily derived from the DILIrank dataset, supplemented with extra non-hepatotoxic compounds collected from seven additional literature sources. As previously described, only compounds consistently classified as non-hepatotoxic in at least two of these datasets were included^20, 21^. The resulting balanced dataset comprised 869 unique drug compounds, including 432 hepatotoxic and 437 non-hepatotoxic compounds. Among these, 500 compounds also had corresponding L1000 gene expression profiles and were used as the modeling set, which was further divided into training and test sets. Within these 500 compounds, 343 were hepatotoxic and 157 were non-hepatotoxic. To construct a balanced training set, an equal number of hepatotoxic and non-hepatotoxic compounds were selected using scaffold-based splitting. Molecular scaffolds for the 500 compounds were identified using the RDKit Murcko Scaffold algorithm, yielding 338 unique scaffolds (**Supplementary Information**). Scaffold-based partitioning of hepatotoxic drugs produced a subset of 157 hepatotoxic compounds combined with 157 non-hepatotoxic compounds for training. A second scaffold split was applied to this balanced training set to allocate 20% of compounds to combine with the remaining 186 hepatotoxic compounds to form the external test set. The final training and test sets consisted of 251 and 249 compounds respectively. This scaffold-based splitting strategy ensured the balance of two classes in the training set and structural diversity of both training and test sets.

Potential MIE assays for the AOP network were retrieved from ToxCast/Tox21 database. After examining the bioassay target information, assays that measure key molecular interactions relevant to toxicity mechanisms were prioritized. MIEs represent the initial interactions between a molecule and a biological system, such as the binding of a chemical to a specific receptor or enzyme inhibition^22, 23^. Thus, we selected 272 assays that were classified under receptor activation, receptor binding, regulation of catalytic activity, or regulation of transporter activity (**Table S1**). Missing data is common when using HTS assay results for modeling target compounds. While QSAR approaches are not always reliable for complex *in vivo* endpoints, they can effectively predict *in vitro* assay responses with simpler, well-defined mechanisms for untested compounds^24^. Among the 272 initially selected assays, those with fewer than 10 experimental active results were excluded, leaving 215 MIE assays for QSAR modeling (**Table S1**). For each assay, the training set consisted of active/inactive compounds balanced by including all compounds from the minority class and randomly sampling an equal number from the majority class. An in-house auto-QSAR workflow was then employed to develop predictive models for these 215 MIE assays. Each assay was modeled using combinations of six machine learning algorithms and four types of chemical descriptors. Five-fold cross-validation was used to train and evaluate the models, and the best-performing model for each assay was selected. As shown in **Fig. 2B,C**, more than 50% of the models achieved balanced accuracy above 0.7. The t-SNE projections of training and test set compounds based on Saagar fingerprints, MIE profiles, and gene expression profiles are shown in **Fig. S1**. Test set compounds occupied a similar feature space to training compounds, and no clear separation between hepatotoxic and non-hepatotoxic compounds was observed.

**Fig. 2.**
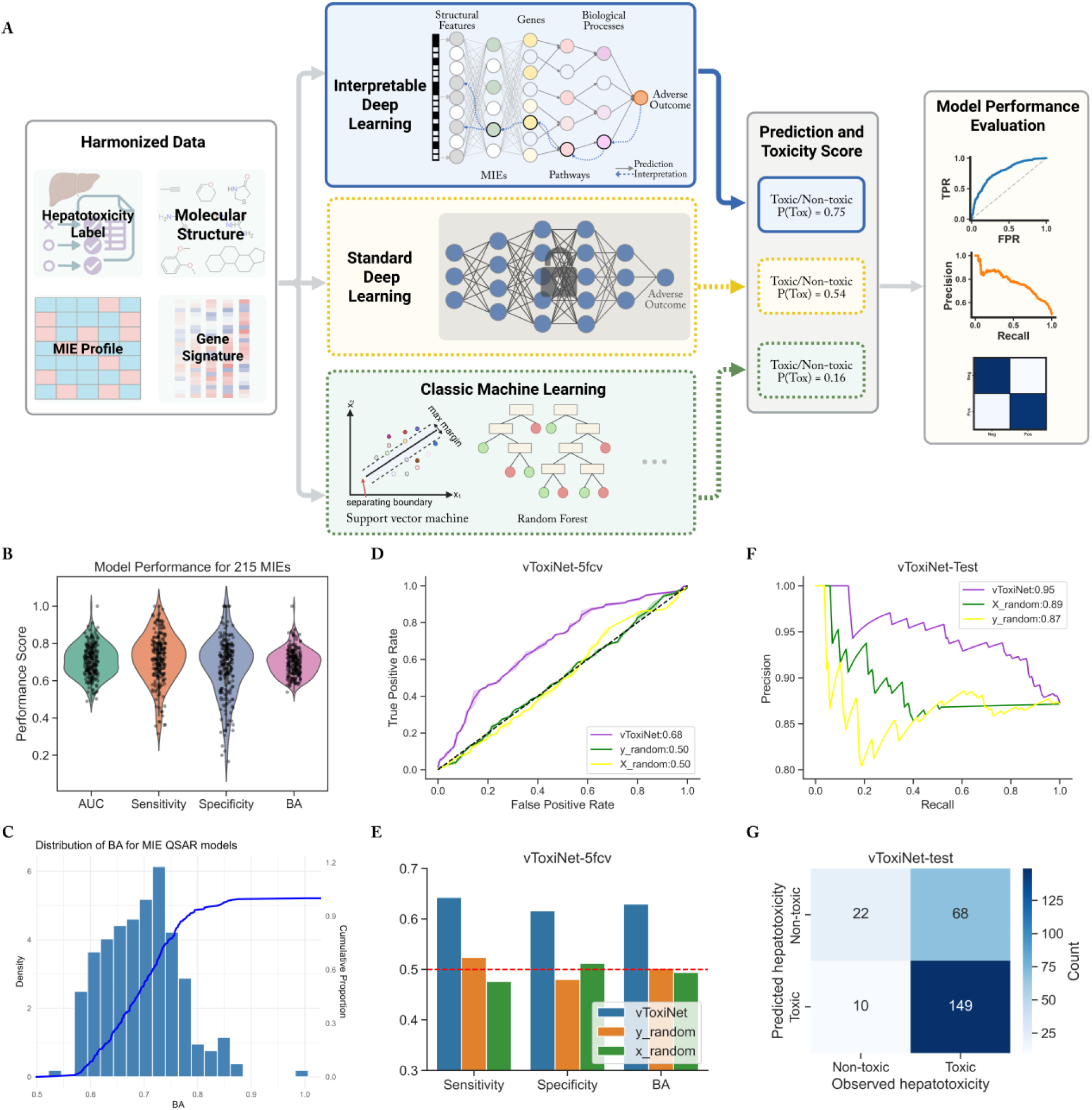
Evaluation of vToxiNet performance and robustness. **(A)** Schematic overview of the vToxiNet performance evaluation workflow. **(B)** Boxplots summarizing the distribution of model performance across MIE assays. **(C)** Histogram and cumulative distribution of balanced accuracy (BA) for models trained on MIE assays. **(D-G)** Performance of vToxiNet and control models. Two control models were constructed to assess robustness: **X_random**, in which the Saagar fingerprint, MIE, and gene expression profiles were randomly permuted; and **y_random**, in which hepatotoxicity labels were randomly permuted. **(D)** Five-fold cross-validation ROC curves and corresponding AUROC values for vToxiNet and control models. **(E)** Classification metrics, including sensitivity, specificity, and balanced accuracy (BA). **(F)** Precision-recall curves and average precision (area under the precision-recall curve) for vToxiNet and control models on the test set (n = 249). **(G)** Confusion matrix of vToxiNet predictions on the test set (n = 249).

### 2.2. Construction and predictive performance of vToxiNet

vToxiNet integrates multimodal data and AOP-based biological knowledge into a neural network modeling framework for hepatotoxicity. A total of 834 Saagar fragments were used as input features, followed by a hidden layer representing 215 MIEs, a hidden layer representing 280 liver injury-related genes, and five subsequent hidden layers encoding 1,119 biological pathways corresponding to potential key events. The network concludes with an output layer representing the adverse outcome, i.e., hepatotoxicity. The complete Reactome hierarchy contains 11,727 genes and 2,751 pathways, arranged in a complex graph with maximum depth of 10^25, 26^. Directly mapping this full hierarchy into neural network framework is computationally infeasible due to the high dimensionality and excessive number of hidden layers. To reduce model complexity and focus on hepatotoxicity-relevant mechanisms, we trimmed the pathway hierarchy to include only genes associated with chemical and drug induced liver injury and their corresponding pathways. The curated hierarchy consisted of 280 genes and 1,119 related pathways organized into five pathway layers (**Fig. S2**, **Supplementary Information)**.

Connections between the gene layer and the first pathway hidden layer were constrained based on gene-pathway annotations, while connections among pathway hidden layers followed established child-parent relationships. Each pathway neuron received input from its child pathway nodes in the preceding layer and transmitted output to its parent pathway in the subsequent layer. This biologically grounded hierarchy produced a sparse and interpretable network structure, in contrast to conventional fully connected neural networks.

vToxiNet was trained using a set of 251 drugs and tested on an external set of 249 drugs with distinct molecular scaffolds. The overall evaluation workflow is illustrated in **Fig. 2A**. Model construction was evaluated using five-fold cross-validation, a standard approach for assessing model performance^15^. Predictions from all test folds were combined to calculate overall performance metrics. The trained vToxiNet model achieved a five-fold cross-validation AUROC of 0.68, sensitivity of 0.64, specificity of 0.62, balanced accuracy of 0.63, and average precision of 0.67 (**Fig. 2D,E, Fig. S3A**). To further confirm model robustness, permutation test was conducted. In the first control network (*X_random*), input Saagar fingerprints, MIE profiles, and gene expression profiles were randomly shuffled before training. Across 100 repetitions, this control model yielded a mean AUROC of 0.49 (SD = 0.05) and mean balanced accuracy of 0.50 (SD = 0.04), consistent with random prediction (**Fig. 2D,E**). In the second control network (*y_random*), hepatotoxicity labels were randomized while keeping the input features unchanged. This model achieved a mean AUROC of 0.51 (SD = 0.03) and mean balanced accuracy of 0.50 (SD = 0.04), further confirming the robustness of the resulting vToxiNet.

We next evaluated the model using the external test set of 249 compounds. The vToxiNet achieved an AUROC value of 0.74 and average precision of 0.95 on the test set (**Fig. 2F, Fig. S3B**). The model correctly predicted 69% (149 out of 217) of hepatotoxic compounds and 69% (22 out of 32) of non-hepatotoxic compounds (**Fig. 2G**), indicating balanced predictivity for both hepatotoxic and non-hepatotoxic compounds.

### 2.3. Performance comparison of vToxiNet with conventional machine learning models

We compared the performance of vToxiNet with conventional machine learning-based QSAR models (**Fig. 3, Fig. S3C-H**). First, a dense neural network with the same number of neurons per layer as vToxiNet, but without incorporating biological data or structural constraints, was trained using Saagar fingerprints as input features. In addition, QSAR models were developed using traditional machine learning algorithms, including random forest (RF) and support vector machine (SVM), with Saagar fingerprints as descriptors (**Fig. 3A-C, Fig. S3C,D**). Additional models were trained using MIE activity profiles or gene expression profiles as input features (**Fig. 3D-H, Fig. S3E-H**). Across five-fold cross-validation and external test evaluation, vToxiNet achieved comparable or superior predictive performance relative to all benchmark models. Models trained only on gene expression profiles exhibited the weakest overall performance, likely due to higher noise and lower signal specificity^27^. Together, these results demonstrate that integrating biological hierarchy with multimodal data enhances model robustness and predictive capacity compared with conventional QSAR approaches.

**Fig. 3.**
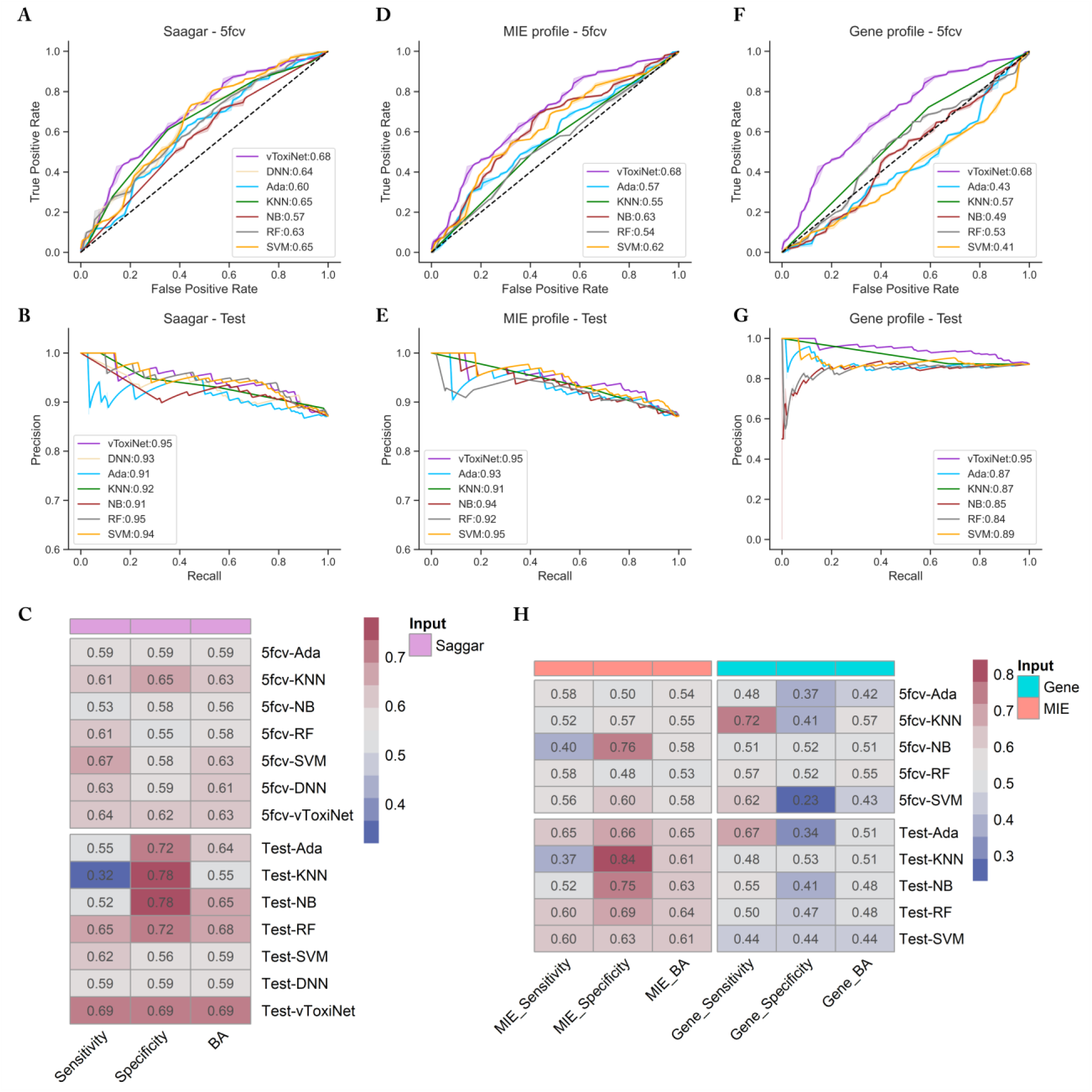
Benchmark comparison of vToxiNet with conventional machine learning models. (A-C) Performance comparison between vToxiNet and classical machine learning models trained using Saagar fingerprints as input features: **(A)** five-fold cross-validation ROC curves, **(B)** test set precision-recall curves, and **(C)** summary classification metrics. **(D-H)** Performance comparison between vToxiNet and classical machine learning models trained using MIE profiles or gene expression profiles as input features: **(D, F)** five-fold cross-validation ROC curves, **(E, G)** test set precision-recall curves, and **(H)** summary classification metrics.

### 2.4. Model interpretation identifies genes and pathways driving hepatotoxicity

This biologically informed architecture of vToxiNet enables mechanistic tracing through the hidden layers. Using the Layer-wise Relevance Propagation (LRP) approach, predictions of target chemicals can be back-propagated through the network to assign relevance (𝑅) scores to individual neurons^28^. These 𝑅 scores quantify each neuron’s contribution to the final output, allowing genes and pathways to be ranked by importance. **Fig. S4** visualizes the hierarchical structure of the gene-pathway network using a sunburst plot, illustrating interpretable key events layers. The 280 DILI-related genes were distributed across multiple biological processes, including metabolism, immune response, and cellular response to stimuli, etc. Among these, metabolism-related pathways involved the largest number of genes (n = 104).

To evaluate the relative importance of specific genes related to hepatotoxicity, we compared both gene 𝑅 scores derived from the vToxiNet model and L1000 ModZ gene expression values between hepatotoxic and non-hepatotoxic compounds. After model training, 𝑅 scores were calculated for each compound in the training set. The distribution of log₂-transformed absolute gene 𝑅 scores for the 251 training compounds across hepatotoxic and non-hepatotoxic groups is shown in **Fig. 4A**, where a large portion of hepatotoxic compounds exhibited higher gene 𝑅 values. A volcano plot was generated to visualize the differences in gene 𝑅 scores between hepatotoxic and non-hepatotoxic groups (**Fig. 4B**), revealing a large number of genes with significantly higher relevance scores in hepatotoxic compounds. The top 15 ranked genes are listed in **Table S2**, several of which are functionally related to xenobiotic metabolism, lipid homeostasis, and hepatic stress responses. For example, cytochrome P450 enzymes *CYP2A6*, *CYP2B6*, and *CYP1A2* are phase I drug-metabolizing enzymes that mediate oxidative biotransformation of many therapeutic compounds^29, 30^ (**Fig. S5, S7**). *GSTM4*, a member of the glutathione S-transferase Mu family, participates in phase II detoxification through glutathione conjugation of reactive metabolites^31^. Genetic polymorphisms or altered expressions of these genes may associate with interindividual variability in drug clearance and susceptibility to hepatotoxicity^32, 33^. Additionally, *FMO3* (flavin-containing monooxygenase 3), plays a role in xenobiotic oxidation, and its dysfunction has been linked to abnormal drug metabolism and endoplasmic reticulum stress^34, 35^. The distributions of L1000 ModZ scores for the 500 modeling compounds were largely overlapping between the hepatotoxic and non-hepatotoxic groups, indicating that most genes did not exhibit significant differential expression between the two groups (**Fig. 4C)**. Volcano plot visualizes the differences in gene expression (ΔModZ) between hepatotoxic and non-hepatotoxic compounds (**Fig. 4D**). One gene, *RBP1* (encodes cellular retinol-binding protein 1, CRBP-1), showed higher expression (*p* < 0.05) in the hepatotoxic group, and four genes *OGDH*, *PANX1*, *CXCL10*, and *PC* showed higher expression (*p* < 0.05) in the non-hepatotoxic group. These genes are involved in essential processes related to metabolism, cellular stress and immune responses. For example, *PC* (pyruvate carboxylase) maintains anaplerotic flux for gluconeogenesis, and its inhibition can predispose liver to oxidative stress and inflammation^36^. Although most genes did not show significant expression differences between the two groups, small changes across multiple genes may collectively influence the pathways and biological processes involved in the adverse outcome process. vToxiNet can capture these subtle shifts as they propagate through the downstream pathway layers.

**Fig. 4.**
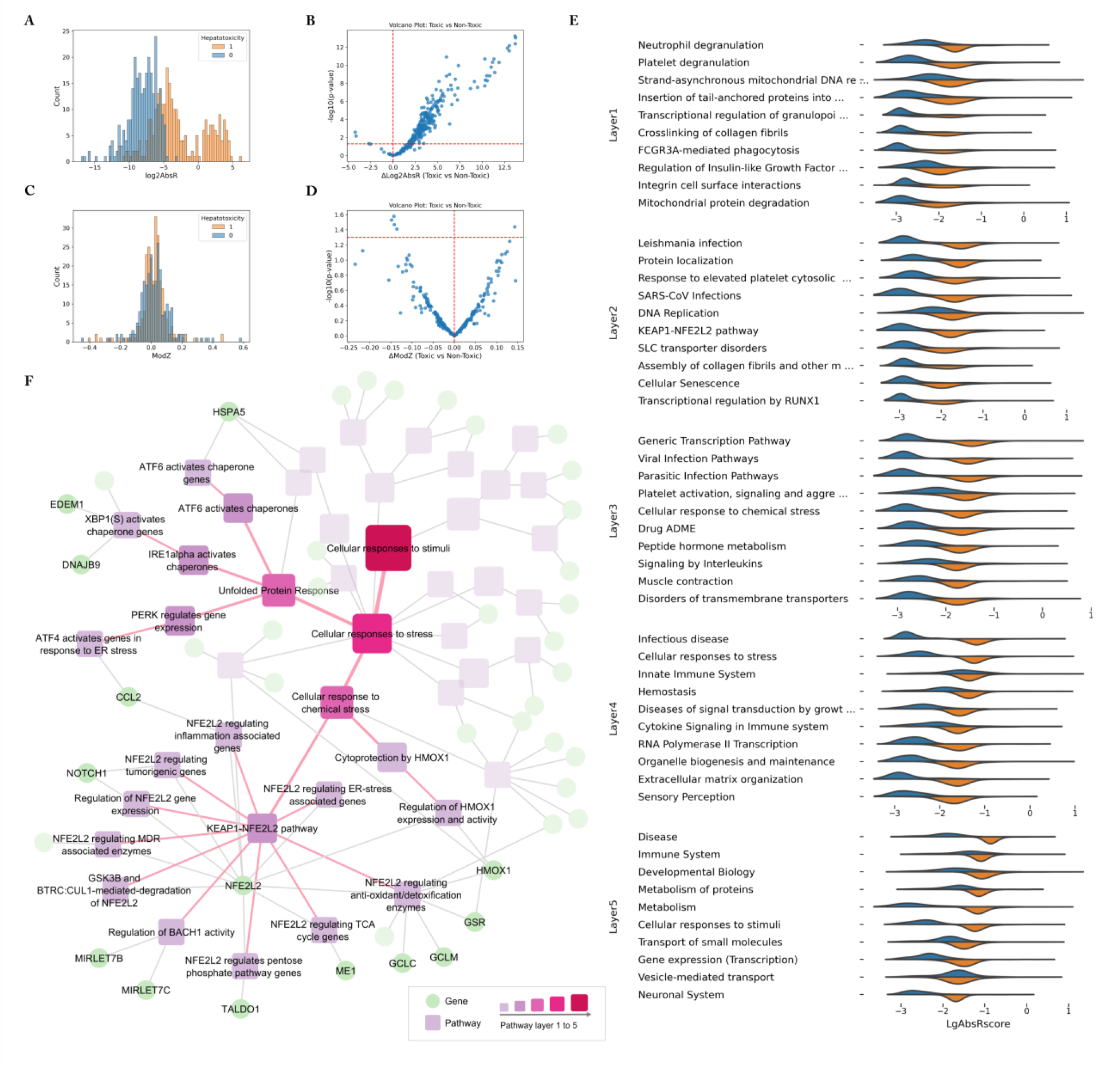
Identification of potential gene and pathway biomarkers for hepatotoxicity. **(A)** Distribution of gene relevance (𝑅) score for hepatotoxic and non-hepatotoxic compounds. **(B)** Volcano plot showing differences in gene relevance (Δlog_2_AbsR) and statistical significance. **(C)** Distribution of differential gene expressions among the hepatotoxic and non-hepatotoxic compounds. **(D)** Volcano plot showing differences in expression (ΔModZ) and statistical significance. Δlog_2_AbsR or ΔModZ are calculated as mean(toxic) − mean(non-toxic). **(E)** Visualization of pathway-level contributions to hepatotoxicity prediction, showing the distribution of R scores for the top 10 pathways in each hierarchical layer. Orange denotes hepatotoxic compounds and blue denotes non-hepatotoxic compounds. **(F)** Gene-pathway hierarchy within the *Cellular responses to stimuli* branch. Only nodes forming a complete hierarchical chain and exhibiting significantly higher log_2_AbsR values in hepatotoxic compounds compared to non-hepatotoxic compounds (FDR < 0.01) are labeled.

Pathway 𝑅 scores were calculated for all nodes in the five key event layers and ranked by their median 𝑅 values among the training compounds. Top 10 ranked pathways in each layer were shown in **Fig. 4E** with the distribution of 𝑅 scores across hepatotoxic and non-hepatotoxic groups. Several pathway nodes exhibited overall higher 𝑅 scores in hepatotoxic samples, and differences between the two groups became more pronounced in higher layers, particularly among highly ranked nodes. For example, in layers 3 and 4, hepatotoxic compounds displayed distinct 𝑅 score distributions compared with non-hepatotoxic compounds. The top-ranked node in layer 4, *Cellular responses to stimuli*, showed a larger difference between the two groups than *Post-translational protein modification*. These results indicate that vToxiNet captures local pathway activations that contribute differently to the overall prediction, with certain biological processes preferentially activated in hepatotoxic compounds^37^. The biological processes branches showing strong differentiation across multiple layers were *Metabolism* (**Fig. S5**), and *Cellular responses to stimuli* (**Fig. S6**), which may represent potential mechanistic pathways underlying chemical and drug induced hepatotoxicity. The liver is particularly vulnerable to toxicity because of its central role in metabolizing and eliminating xenobiotics, especially lipophilic drugs^2^. This process is controlled by large families of proteins and enzymes, such as those represented in **Fig. S5**, which collectively influence the accumulation of exposure to drugs and their metabolites.

Metabolism of the xenobiotics and drugs can produce reactive metabolites that triggers oxidative stress responses, an important component of *Cellular responses to stimuli* process (**Fig. S6**). In particular, the KEAP1-NFE2L2 signaling pathway plays a pivotal role in maintaining redox homeostasis and protecting hepatocytes from xenobiotic-induced oxidative stress (**Fig. 4F**).

Upon exposure to reactive oxygen species (ROS) or electrophilic metabolites, KEAP1 cysteine residues are oxidized, allowing NFE2L2 to accumulate and translocate to the nucleus, where it activates transcription of antioxidant genes such as *GSTM4*, *GCLC* (Glutamate-cysteine ligase), *NQO1* (NAD(P)H quinone dehydrogenase 1)^38, 39^. Impairment or inadequate of this adaptive response through genetic variation, or sustained oxidative overload, can cause hepatocyte death and heighten susceptibility to drug-induced liver injury^2^. In contrast, the distributions of 𝑅 scores for individual Saagar fragments or MIE nodes showed minimal differences between hepatotoxic and non-hepatotoxic groups (**Fig. S8**), likely reflecting the multifactorial nature of hepatotoxicity, which arises from the combined perturbation of multiple pathways rather than from the alteration of a single molecular target.

### 2.5 Model validation using independent external datasets

We further evaluated vToxiNet using an independent external hepatotoxicity dataset. This dataset included compounds with known hepatotoxicity classifications from DILIst^40^, the NCTR Liver Toxicity Knowledge Base (LTKB) benchmark dataset^41^, Greene’s data set^42^, and Xu’s data set^43^. Compounds overlapping with the modeling set were removed, yielding 267 unique compounds (165 hepatotoxic and 92 non-hepatotoxic) with available L1000 gene expression data. Missing MIE assay results were imputed using the pre-trained QSAR models. The external validation set contained 206 unique Murcko molecular scaffolds, 30 of which were not present in the training set. vToxiNet achieved an AUROC of 0.61 on the independent external validation set, and compounds with prediction probabilities ≥ 0.88 demonstrated a precision of 0.81, reflecting strong enrichment in the high-confidence predictions (**Fig. S9**).

In **Fig. 5A**, we highlight five scaffolds with high predictive accuracy, along with representative compounds containing those scaffolds. Model interpretation revealed that compounds sharing the same scaffold can exhibit similar mechanistic signatures. For example, the first scaffold represents a steroid core. Among the four compounds containing this scaffold, three were correctly predicted as true negatives, whereas nandrolone (PubChem CID 9904) was misclassified as a false negative. According to LiverTox® and previous reports, the mechanism of nandrolone-induced liver injury is not well defined but may involve alterations in immune function and typically occurs only with prolonged use or high doses^44^. Such exposure-dependent scenarios cannot be fully captured by the current vToxiNet model, which is designed to characterize intrinsic toxicity responses based on chemical structure and biological perturbations. Analysis of the top contributing pathways for these four compounds revealed consistent patterns, with *Immune system* processes showing the highest relevance and limited involvement of metabolic pathways (**Fig. S10**).

**Fig. 5.**
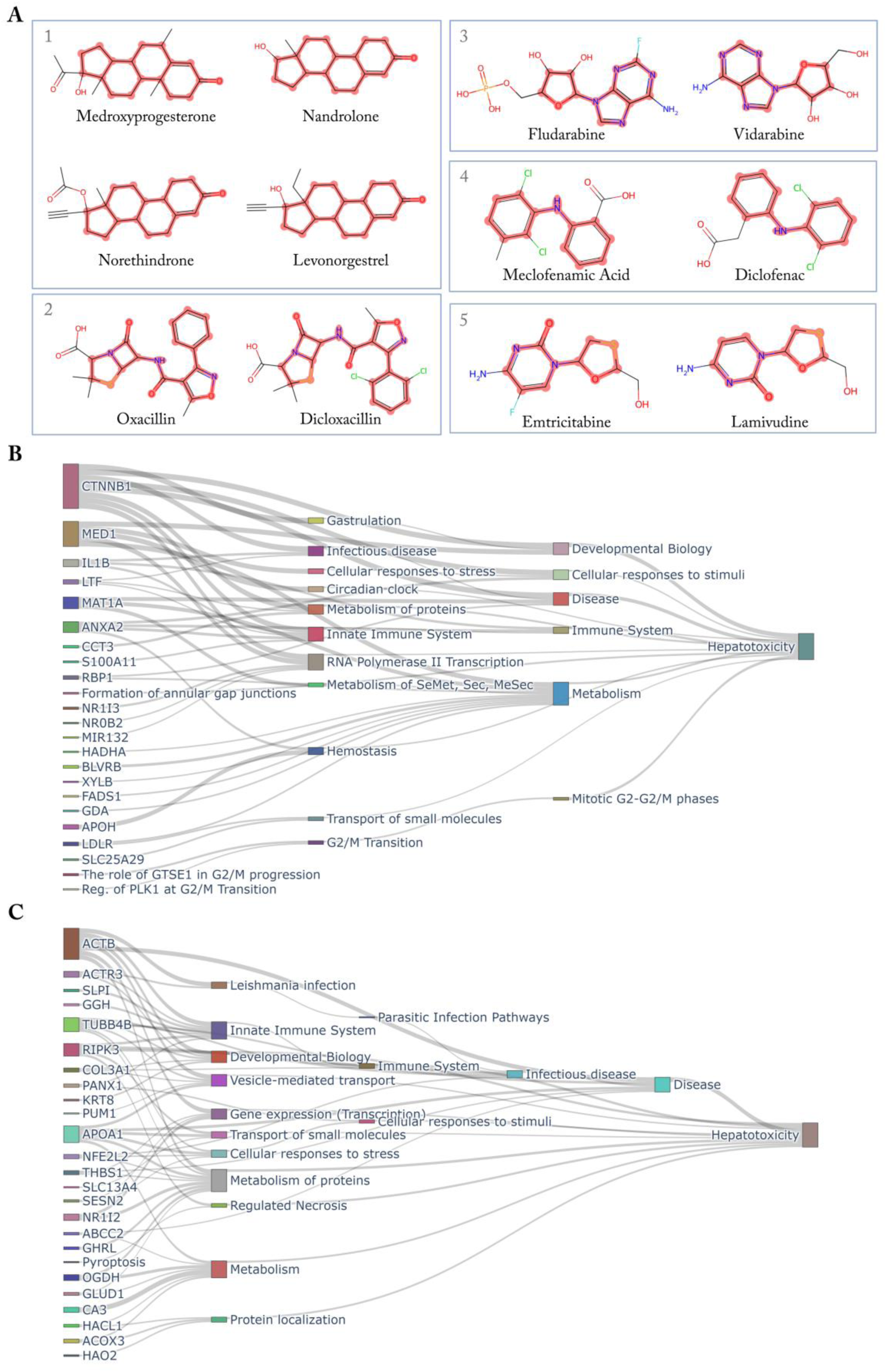
Model predictions for compounds in the independent external dataset. **(A)** Top-ranked scaffolds and their representative drug compounds. Scaffolds were ranked by the proportion of correctly predicted compounds among all compounds containing each scaffold. Notably, scaffolds 2 and 5 were not present in the training set. (**B-C**) Visualizations of the top ranked genes and pathways with high contributions to hepatotoxicity predictions (𝑅 score) for diclofenac **(B)** and meclofenamic acid **(C)**.

Diclofenac (CID 3033) and meclofenamic acid (CID 4037) both contain a diphenylamine scaffold and were correctly predicted as true positives. The mechanism of diclofenac-induced hepatotoxicity is multifactorial, involving both immune-mediated effects and the formation of reactive metabolites that can cause mitochondrial damage^45–47^. These findings are consistent with the top-ranked genes and pathways identified by vToxiNet, in which nodes related to metabolic regulation (e.g., *MED1*) and mitochondrial function (e.g., *MAT1A*) showed higher contribution to the model’s hepatotoxicity predictions (**Fig. 5B**, **Fig. S10**). The mechanism of meclofenamic acid-induced hepatotoxicity is less clearly understood but likely involves metabolic bioactivation to reactive intermediates that trigger immune responses^48^. Consistent with this, vToxiNet ranked *Innate immune system* and *Cellular response to stress* among the top pathways, with multiple related genes showing elevated 𝑅 scores (**Fig. 5C**, **Fig. S10**). Collectively, these results demonstrate that vToxiNet not only maintains predictive accuracy across novel chemical but also captures dominant activation of potential key events, enabling comparison and inference of underlying mechanistic patterns.

## Discussion

Understanding and predicting complex biological outcomes such as drug-induced hepatotoxicity remain challenging due to the multiscale organization of biological systems, in which molecular perturbations propagate across genes, pathways, and higher-order cellular processes. Although recent advances in machine learning have improved predictive performance for toxicity endpoints, most models treat biological measurements as unstructured features, limiting interpretability and obscuring links between predictions and underlying mechanisms.

Here, we developed vToxiNet, an interpretable deep learning framework that integrates chemical descriptors, assay responses, transcriptomic profiles, and the Reactome pathway hierarchy within a biologically constrained architecture. By embedding molecular initiating events, gene-level responses, and pathway structure into a unified network, vToxiNet operationalizes adverse outcome pathway concepts while enabling mechanistic interpretation alongside prediction.

Rather than relying solely on post hoc explanation, this design allows mechanistic signals to be traced across biological scales, linking chemical perturbations to pathway-level processes associated with liver injury.

Overall, vToxiNet demonstrates that integrating biological hierarchy with multimodal data can enhance both the accuracy and interpretability of hepatotoxicity prediction, providing a foundation for transparent and mechanism-informed NAMs in predictive toxicology. Several limitations should be noted. While the Reactome hierarchy provides biologically grounded structure, it encompasses broad cellular processes that are not toxicity-specific, which may dilute hepatotoxicity-relevant signals. In addition, hepatotoxicity classification criteria vary across different resources and exposure context, particularly for cases involving misuse or long-term exposure, creating challenges for model training and evaluation. Future integration of high-quality exposure and pharmacokinetic data may further strengthen predictive robustness and translational relevance. Collectively, these findings support biologically constrained deep learning as an appliable strategy for integrating structured biological knowledge into interpretable models of complex toxicological outcomes.

## 3. Methods

### 3.1. Hepatotoxicity Dataset

Hepatotoxicity labels were collected from the DILIrank dataset and other literatures. DILIrank dataset contains 1036 FDA-approved drugs that are divided into four classes: three classes (Most-, Less-, No-DILI concern) with confirmed causal evidence linking drugs to potential/no liver injury, and one additional class (Ambiguous-DILI-concern) with causality undetermined^49^. Compounds belonging to the Most or Less-DILI-concern were classified as *hepatotoxic*. Compounds designated as No-DILI-concern were classified as *non-hepatotoxic*. Additional compounds were collected from seven hepatotoxicity datasets in literatures that were used in our previous studies^20, 21, 50^. The standards for harmonization of various hepatotoxicity classifications into binary categories of 1 (hepatotoxic) and 0 (non-hepatotoxic) were described previously^50^. Compounds with conflicting classifications across the DILIrank and seven literature datasets were removed. Only compounds with consistent classifications (i.e., have at least two consistent hepatotoxic classifications among these datasets) were selected.

### 3.2. ToxCast/Tox21 HTS Data

The ToxCast and Tox21 projects generated large-scale HTS data by testing ∼10,000 of environmental and pharmaceutical compounds across thousands of human cell–based assays, enabling mechanistic interpretation of toxicity pathways^51, 52^. For this study, the public HTS bioassay hit calls and assay endpoint information were downloaded from the EPA ToxCast/Tox21 database (Invitrodb 3.4). Assay metadata included assay design type, target gene, and biological process targets etc. Assays that can be mapped as molecular initiating events (MIEs) in the AOP framework were selected for model training, specifically those targeting receptor activation, receptor binding, regulation of catalytic activity, or regulation of transporter activity. Corresponding chemical structures, in the form of SMILES strings, were obtained using the NCI/CADD Chemical Identifier Resolver (https://cactus.nci.nih.gov/) using CAS numbers.

### 3.3. L1000 Gene Expression Data

The Connectivity Map and its next-generation L1000 platform generated large-scale transcriptomic profiles by treating human cell lines with ∼20,000 small-molecule perturbagens, using a cost-effective, high-throughput transcriptomic assay that measures 978 ‘landmark’ genes and infers ∼81% of the remaining transcriptome^53^. The most frequently tested cell lines were MCF7 (human breast cancer) and PC3 (human prostate cancer). L1000 data are organized into five levels, with Level 5 representing replicate-collapsed Z-scores (ModZ) that indicate changes in gene expression relative to DMSO-treated controls. For this study, Level 5 data from L1000 was downloaded from the Gene Expression Omnibus (accession numbers GSE92742 and GSE70138). Whenever available, experiments conducted in HepG2 (liver carcinoma) cells were used; otherwise, MCF7 data were selected. Transcriptomic signatures tested at a 24-hour treatment duration were chosen. For chemicals tested at multiple concentrations, the 10 μM condition was selected. When multiple profiles existed for the same chemical, cell line, duration, and concentration, the ‘exemplar’ profile designated by the LINCS consortium was chosen; if no exemplar was available, the profile with the highest ‘tas’ value (transcriptional activity score) was selected.

### 3.4. QSAR Modeling

Not all compounds had been tested in the selected MIE assays, making it necessary to fill data gaps before AOP modeling. Assays suitable for QSAR model development were selected based on having more than 10 active compounds and at least 20 total tested compounds, thereby ensuring a sufficient sample size for reliable model training. For each assay, training set was balanced to include equal numbers of active and inactive compounds. Our in-house auto-QSAR workflow was applied to impute missing values in the MIE assays for target compounds^54^. Six ML algorithms implemented by scikit-learn were employed for modeling: Random Forest (RF), k-Nearest Neighbors (kNN), Support Vector Machine (SVM), Gaussian Naive Bayes (NB), Bernoulli Naive Bayes (BNB), and Adaptive Boosting (AdaBoost). Four types of chemical descriptors were used including extended connectivity fingerprints (ECFP), functional-class fingerprints (FCFP), Molecular ACCess system (MACCS) keys, and RDKit molecular descriptors (www.rdkit.org). Individual models were developed using the combination of one type of descriptor and one of the ML algorithms, resulting in 20 individual models for each assay endpoint. All models were built and evaluated using a standard five-fold cross-validation procedure, with 20% of the training compounds left out for testing purposes during each iteration. For each assay, the best model with highest balanced accuracy was retained to generate predictions.

### 3.5. vToxiNet Model Architecture

The vToxiNet model architecture was constructed using an AOP knowledge-based network framework, beginning with chemical fragments as the input layer, followed by seven hidden layers representing MIEs and potential key events. The network concluded with an output layer that predicts the adverse outcome as hepatotoxicity (**Fig. 1A**).

Saagar fragments were used to support interpretable predictive modeling and read-across in chemical toxicity studies^55^. Saagar fingerprints, represented as binary vectors (1 = presence, 0 = absence) of 834 chemical fragments, were calculated for each modeling set compound and used as inputs to the vToxiNet modeling. The vToxiNet output layer employed a sigmoid activation function, which converts the layer input to a value between 0 and 1 that represents the probability of a compound being hepatotoxic.

Seven hidden layers connected to the vToxiNet input and output layers, organized to represent MIEs and key events within the AOP network. Each layer corresponded to a level of biological responses in AOP. The first hidden layer (MIE layer) incorporated MIE assay outcomes, with each neuron representing one bioassay, as previously described^56^. The second hidden layer (gene layer) incorporated gene expression changes, with each neuron representing one gene associated with chemical and drug induced liver injury. The subsequent five hidden layers were mapped to the Reactome pathway hierarchy filtered by these genes. Curated gene–disease associations (457 genes linked to chemical and drug induced liver injury) were retrieved from the Comparative Toxicogenomics Database^57^ (CTD; MDI Biological Laboratory, Salisbury Cove, ME, and NC State University, Raleigh, NC; https://ctdbase.org/; accessed April 2025).

The complete Reactome pathway hierarchy includes 2,751 human pathways and 11,727 unique genes, with gene-pathway and pathway-pathway connections organized according to child-parent relationships (accessed April 2025)^58^. To reduce dimensionality and extract a hepatotoxicity-relevant pathway hierarchy, we trimmed the Reactome network to only include genes associated with chemical and drug induced liver injury in CTD. Specifically, 280 genes overlapped between Reactome and the CTD chemical and drug induced liver injury gene set. Redundant pathways were pruned, and their child nodes were reconnected to their respective parent nodes, yielding a refined hierarchy containing 280 genes and 1,119 associated pathways organized into five layers. Gene expression change data was derived from L1000 Level 5 ModZ score, where absolute ModZ < 1.5 were classified as unchanged (0), and absolute ModZ ≥ 1.5 were classified as changed (1, upregulated or downregulated).

The input layer, MIE layer and gene layer were densely connected, whereas the connections between gene-pathway and child-parent pathway layers are customized to their biological relationships, resulting in a relatively sparse network^59^. The five pathway hidden layers did not incorporate direct biological activity data and instead used the ReLU activation function to transform the inputs (equation [1] and [2]). The ReLU activation function outputs the input directly if it is positive, otherwise, it outputs zero.

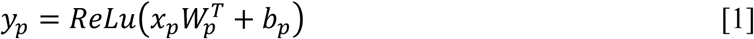

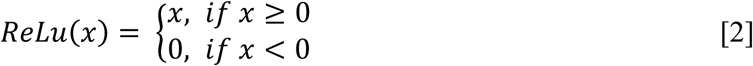

Where 𝑊_𝑝_, 𝑏_𝑝_ are the weight matrix and bias for pathway neuron 𝑝. 𝑥_𝑝_ is the input vector and 𝑦_𝑝_ is the output vector of pathway neuron 𝑝.

Model performance was optimized using binary cross-entropy, a classic loss function for binary classification models^60^. To prevent gradient vanishing and improve the learning of pathway-specific patterns, we adopted the auxiliary layer approach from DCell^61^. Specifically, the output of each pathway module was transformed into an auxiliary scaler 𝑦^_𝑝_ using the Sigmoid function^62^. The auxiliary scalers from all pathway modules were then combined and included the in final binary cross-entropy loss function along with the final output (equation [3]):

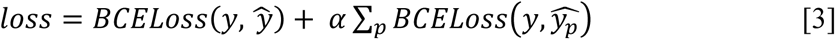

Here, 𝑦 is the observed hepatotoxicity label and 𝑦^ is the final output prediction, 𝛼 is a tunable factor (0 to 1) used during training to balance the contributions from the final output and the individual pathway outputs^61^.

### 3.6. vToxiNet Model Training and Evaluation

The network connection weights were initialized as 0.10 and optimized during training using the Adaptive Moment Estimation (Adam) optimizer^63^ with a batch size of 32 and a learning rate of 0.001. To prevent overfitting, dropout was applied after the fully connected MIE and gene layers, randomly removing 20% of the connections during training^64^. Model training was performed using five-fold cross-validation. In each iteration, four folds were used for training, and the remaining fold was held out for testing. Within the training four folds, 10% of the compounds were further set aside as a validation set for early stopping. Model performance was monitored using balanced accuracy on the validation set after each epoch, and training was stopped if the validation balanced accuracy did not improve for 20 consecutive epochs. Training was terminated after 150 epochs or the early stopping criterion was met. This procedure was repeated until each fold had served once as the test set. Predictions from all test folds were then combined, and final performance metrics, including BA, specificity, precision, sensitivity (recall), (equation [4, 5, 6, 7]), and area under the receiver operating characteristic (ROC) curve (AUROC), were then calculated.

The resulting vToxiNet model outputs a probability score representing the likelihood that a compound is hepatotoxic in humans. A compound was classified as hepatotoxic when the probability exceeded a defined threshold. ROC curves were generated by plotting the true positive rate (TPR) against the false positive rate (FPR) (equation [7, 8]) across varying probability thresholds for distinguishing hepatotoxic from non-hepatotoxic compounds.

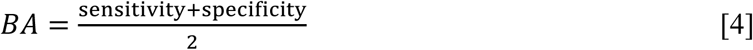

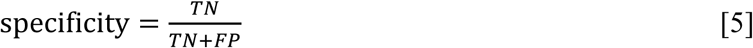

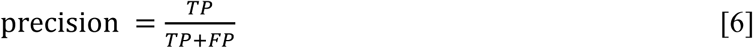

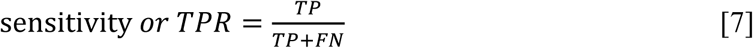

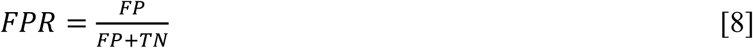

Where TP represents the number of true positives; FP represents the number of false positives; TN represents the number of true negatives; FN represents the number of false negatives.

### 3.7. vToxiNet Model Interpretation

Layer-wise Relevance Propagation (LRP) is an interpretation method for deep neural networks that helps identify the most important neurons contributing to a classification result^65^. LRP leverages the network weights and neural activations from the forward pass to propagate the output backward through the network. In this process, the contribution of each input or intermediate neuron (e.g., representing genes or pathways) can be systematically quantified by a relevance score (*R*). The ε-rule for calculating the *R* score of neuron *j* in layer *l* is given in equation [9, 10].

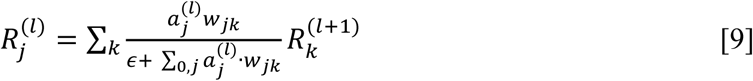

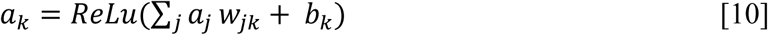

Where 𝑤_𝑗𝑘_ is the weight between neuron 𝑗 and 𝑘, 𝑅 is the 𝑅 score of neuron 𝑘 in the layer 𝑙 + 1, 𝑎_𝑗,_𝑎_𝑘_ are the activations (i.e., output value) of neuron 𝑗 and neuron 𝑘, respectively. 𝜖 is a small constant added to the denominator to make the fraction more stable.

## Data Availability

The data supporting the findings of this study were obtained and curated from publicly available resources and are provided in the Supplementary Information.

## Acknowledgements

This work was supported by the National Institute of Environmental Health Sciences [Grants R01ES031080, R35ES031709, P30ES005022].

## Author contributions

X.J. and T.W. designed and developed the methodology, curated the data, performed formal analysis and investigation, developed the code, conducted validation, and wrote the original draft of the manuscript. D.P.R. contributed to code development. L.M.A. and S.X. provided resources and contributed to manuscript review and editing. H.Z. conceptualized and supervised the study, acquired funding, and contributed to manuscript review and editing. All authors revised the manuscript and approved the final version of the manuscript.

