## Supplementary material for "vToxiNet: a biologically constrained deep learning framework for interpretable prediction of drug-induced hepatotoxicity": Fig. S1-Fig. S10, Table S1-Table S2

[*] Corresponding author

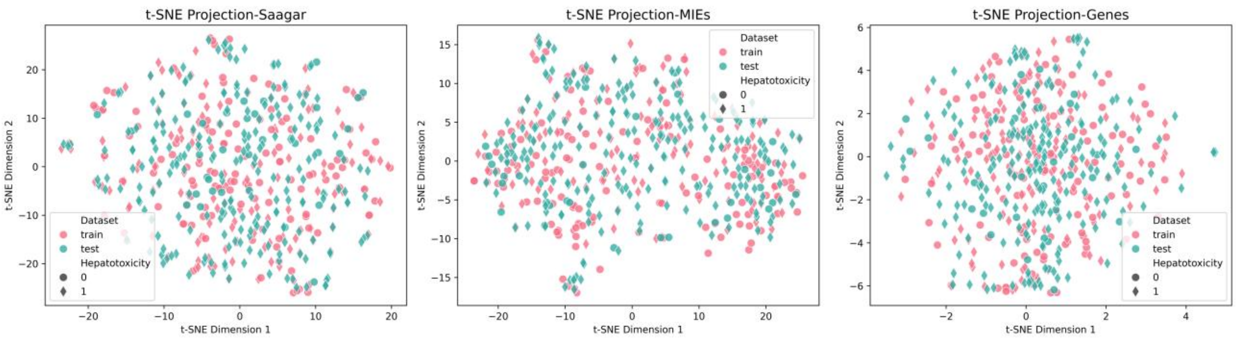


**Fig. S1. t-SNE projections of training and test set compounds using Saagar, MIE profiles or gene expression profiles.**

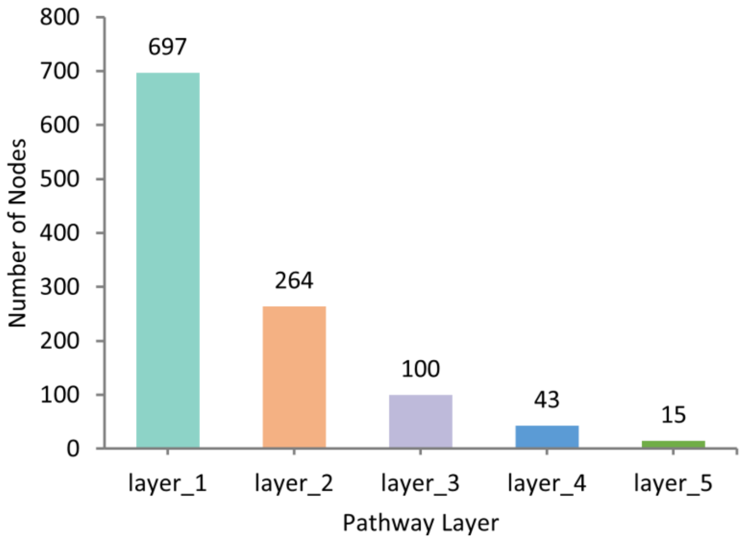


**Fig. S2**. **Number of nodes in each of the pathway layers.**

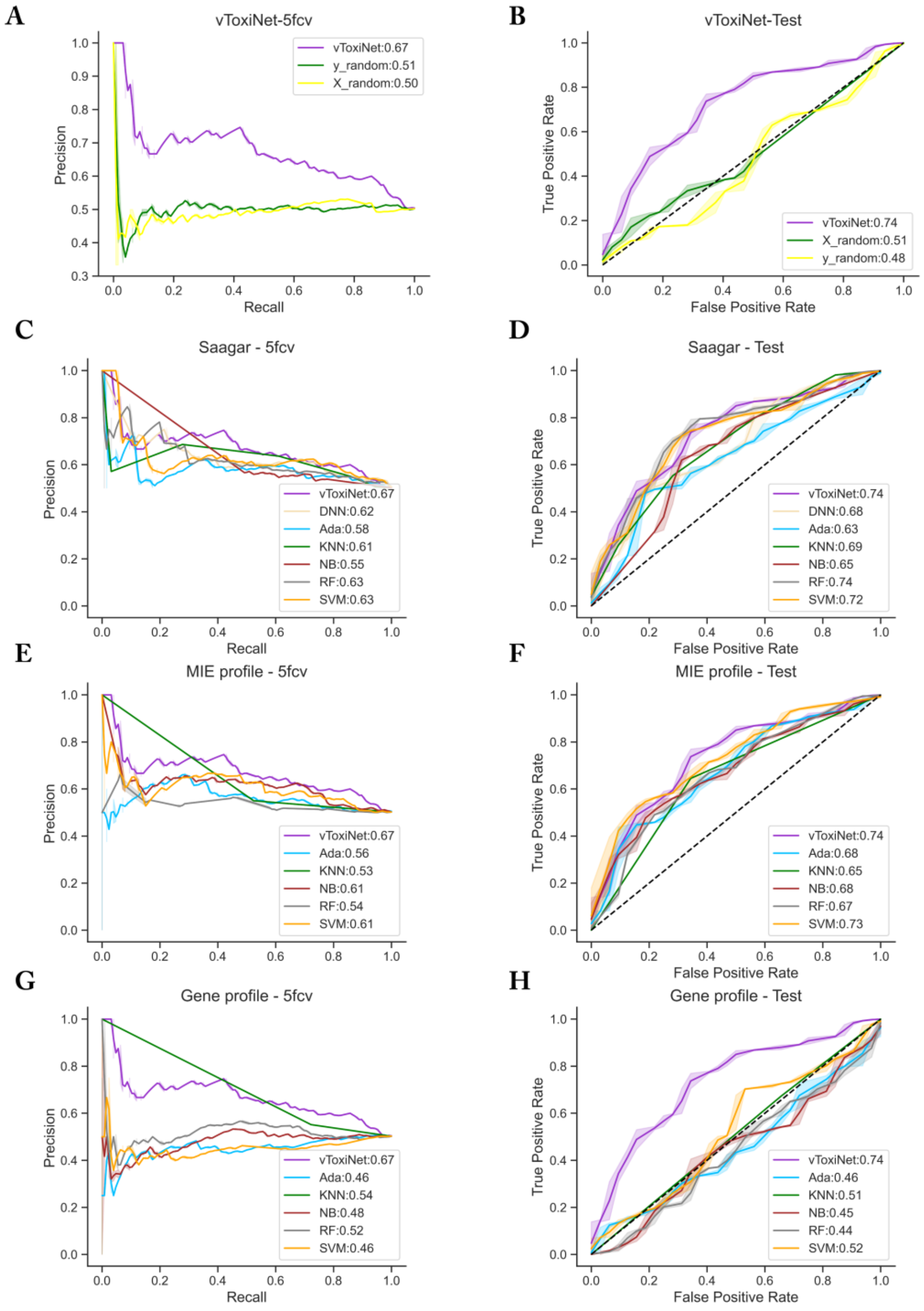


**Fig. S3. Extended evaluation and benchmarking of vToxiNet.** (A, B) Performance of vToxiNet and robustness control models: **(A)** five-fold cross-validation precision-recall curves and **(B)** test-set ROC curves. **(C, D)** Performance comparison between vToxiNet and classical machine learning models trained using Saagar fingerprints as input features: **(C)** five-fold cross-validation precision-recall curves and **(D)** test-set ROC curves. **(E–H)** Performance comparison between vToxiNet and classical machine learning models trained using MIE profiles or gene expression profiles as input features: **(E, F)** five-fold cross-validation precision-recall curves and **(G, H)** test-set ROC curves.

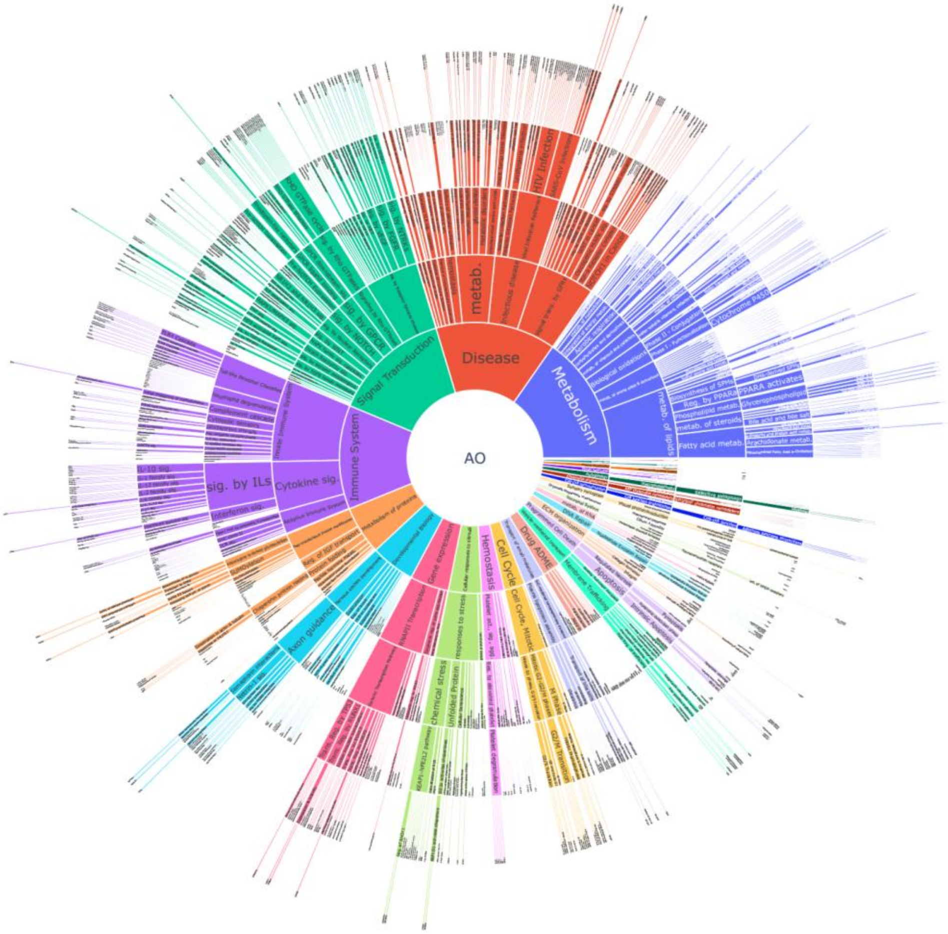


**Fig. S4**. **Sunburst plot visualization of hierarchical structure of the gene-pathway network.**

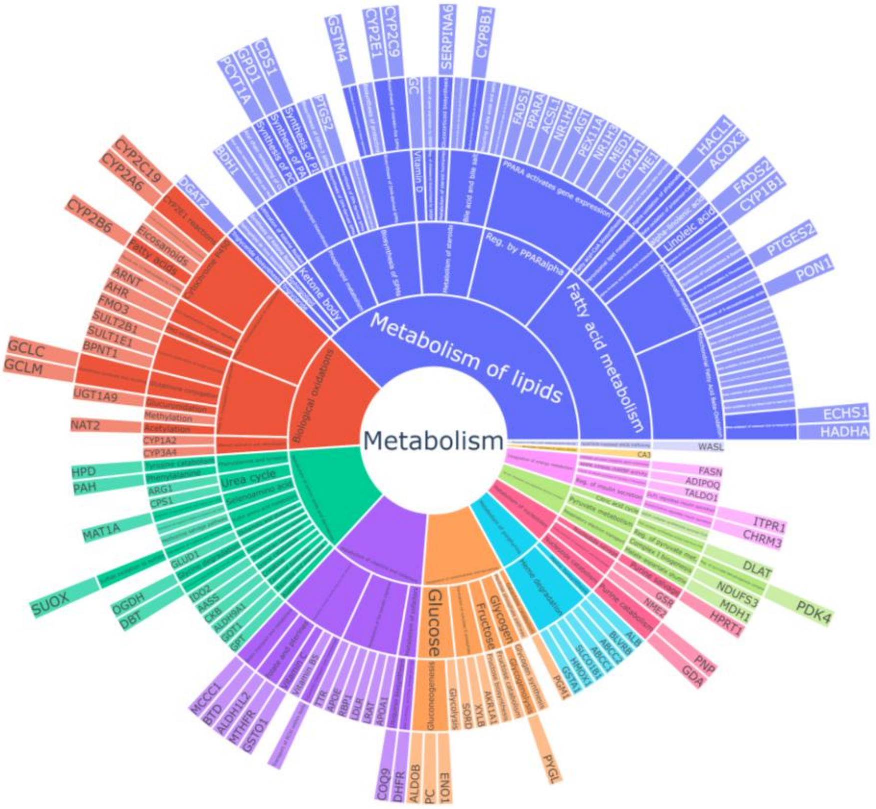


**Fig. S5. Sunburst plot showing the hierarchy of genes and pathways involved in the *Metabolism* process.**

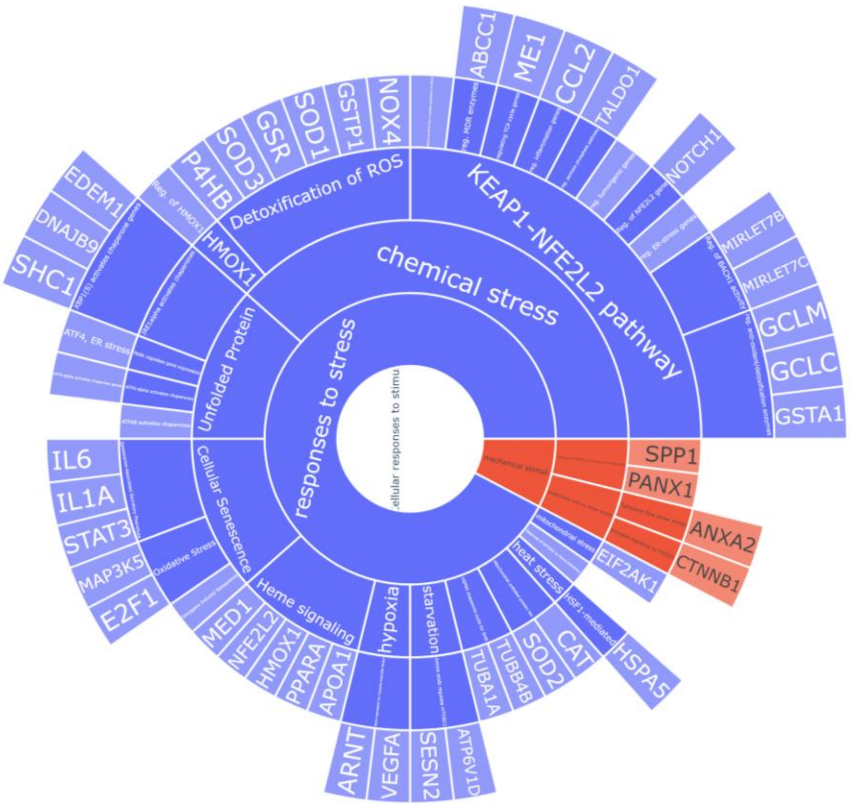


**Fig. S6. Sunburst plot showing the hierarchy of genes and pathways involved in the *Cellular responses to stimuli* process.**

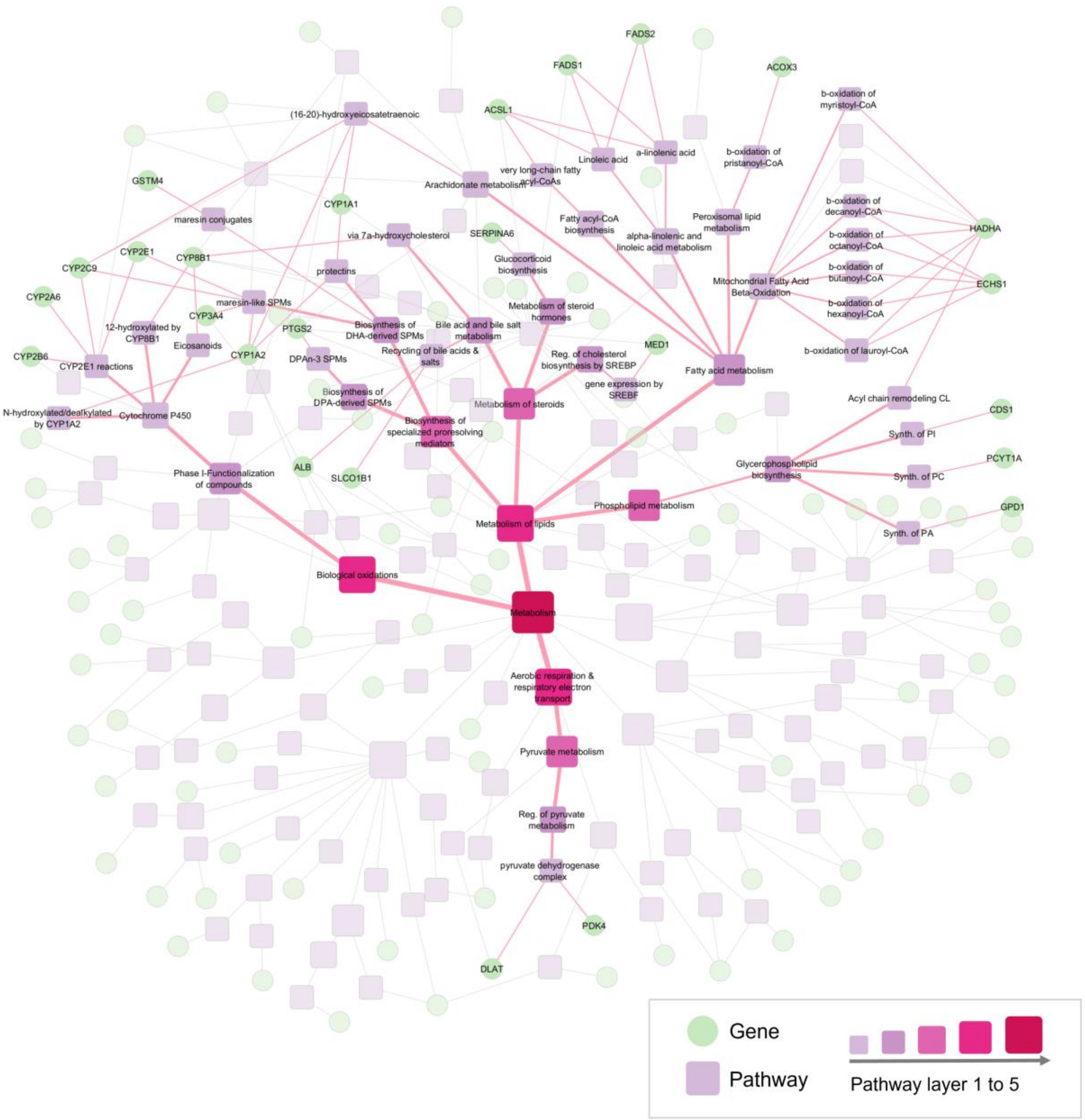


**Fig. S7. Gene-pathway hierarchy within the *Metabolism* branch.** Only nodes forming a complete hierarchical chain and exhibiting significantly higher $log_2$AbsR values in hepatotoxic compounds compared to non-hepatotoxic compounds (FDR < 0.01) are labeled.

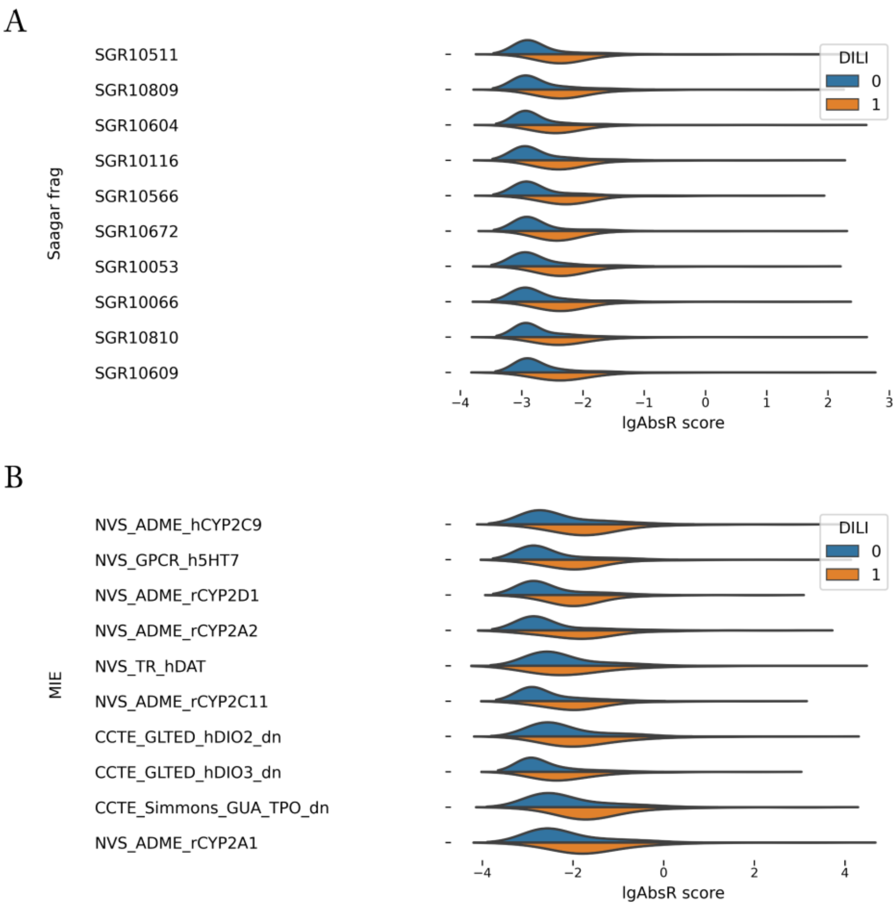


**Fig. S8**. Distribution of *R* scores for top ranked nodes in the **(A)** input layer and **(B)** MIE layer.

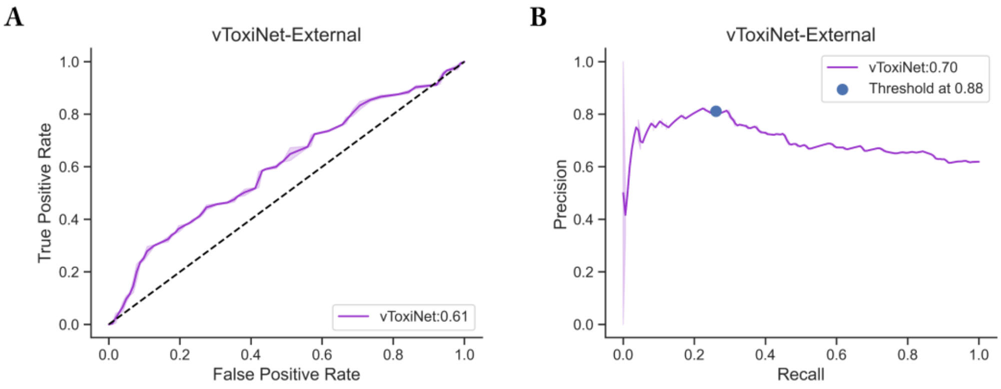


**Fig. S9. Performance of vToxiNet on the independent external dataset.** (A) Receiver operating characteristic (ROC) curve with corresponding AUROC value. (B) Precision-recall curve with corresponding average precision. The blue dot indicates the precision and recall values at a probability threshold of 0.88.

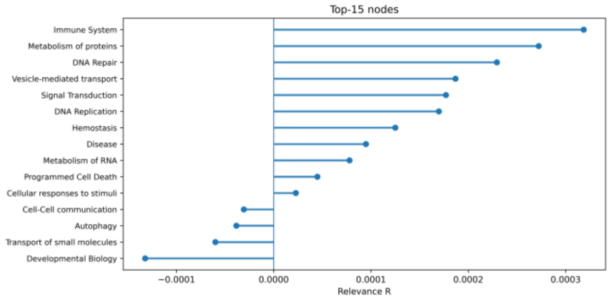


Medroxyprogesterone

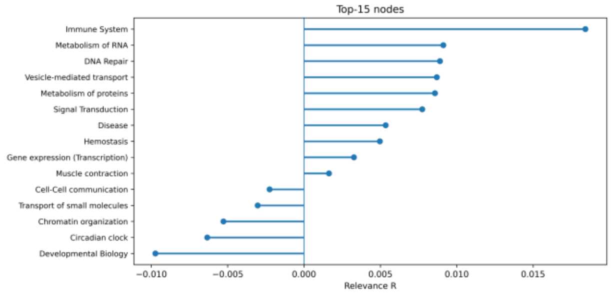


Nandrolone

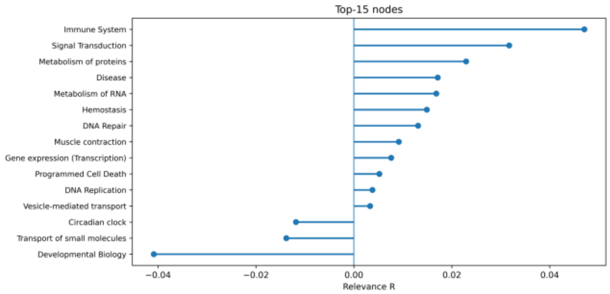


Norethindrone

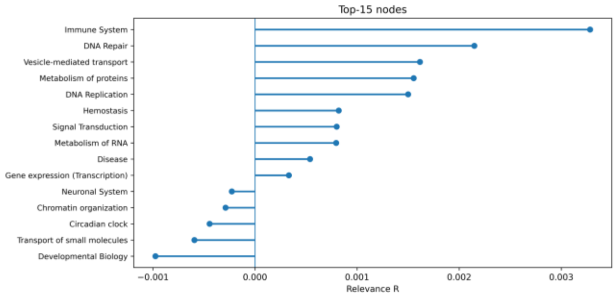


Levonorgestrel

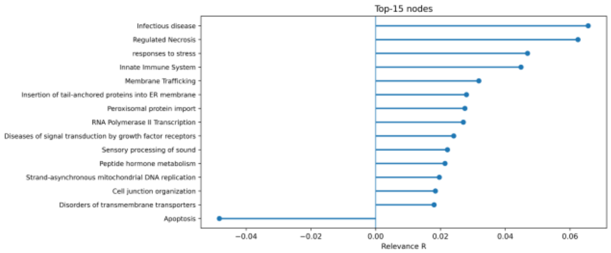


Meclofenamic acid

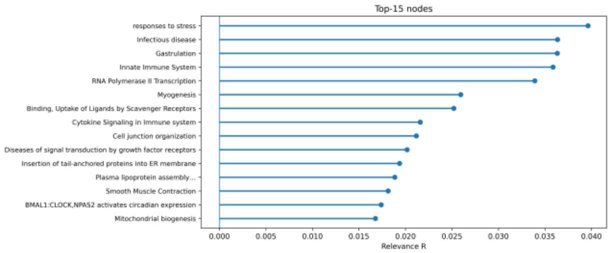


Diclofenac

**Fig. S10**. ***R* scores of top ranked pathways for compounds in the independent external dataset.**

**Table S1**. Candidate MIE assay categories from the ToxCast/Tox21 HTS program.

| biological_process_target | Count | Count QSAR modeling |
|---|---|---|
| receptor activation | 2 | 2 |
| receptor binding | 123 | 109 |
| regulation of catalytic activity | 146 | 103 |
| regulation of transporter activity | 1 | 1 |
| total | 272 | 215 |

**Table S2.** Top 15 Genes that have higher relevance scores in hepatotoxic compounds compared to non-hepatotoxic compounds.

| Gene | Protein name | FDR | Associated Pathway |
|---|---|---|---|
| CYP2A6 | CYP450 family 2 subfamily A member 6 | 1.21E-11 | CYP2E1 reactions |
| GSTM4 | glutathione S-transferase mu 4 | 1.21E-11 | Biosynthesis of maresin conjugates in tissue regeneration |
| PCYT1A | phosphate cytidylyltransferase 1A, choline | 3.74E-11 | Synthesis of PC |
| CYP2B6 | CYP450 family 2 subfamily B member 6 | 4.67E-11 | CYP2E1 reactions |
| CDS1 | CDP-diacylglycerol synthase 1 | 6.87E-11 | Synthesis of PI |
| CYP1A2 | CYP450 family 1 subfamily A member 2 | 1.15E-10 | Aromatic amines be N-hydroxylated or N-dealkylated by CYP1A2 |
| FMO3 | flavin containing dimethylaniline monoxygenase 3 | 4.72E-10 | FMO oxidizes nucleophiles |
| MIRLET7B | microRNA let-7b | 6.06E-10 | Regulation of BACH1 activity |
| PEX11A | peroxisomal biogenesis factor 11 alpha | 6.06E-10 | MLL4 and MLL3 complexes regulate expression of PPARG target genes in adipogenesis and hepatic steatosis |
| FADS2 | fatty acid desaturase 2 | 1.21E-09 | Linoleic acid (LA) metabolism |
| BRD4 | bromodomain containing 4 | 1.21E-09 | Potential therapeutics for SARS |
| NR1I2 | pregnane X receptor (PXR) | 2.49E-09 | Nuclear Receptor transcription pathway |
| GANAB | glucosidase II alpha subunit | 3.43E-09 | Maturation of spike protein |
| GC | GC vitamin D binding protein | 1.26E-08 | Vitamin D metabolism |
| VWF | von Willebrand factor | 1.86E-08 | MAP2K and MAPK activation |

FDR-adjusted p-values were calculated using the Benjamini-Hochberg method
